# Mitochondria tether to Focal Adhesions during cell migration and regulate their size

**DOI:** 10.1101/827998

**Authors:** Redaet Daniel, Abebech Mengeta, Patricia Bilodeau, Jonathan M Lee

**Author notes:** **Corresponding Author:** Jonathan M Lee. Department of Biochemistry, Microbiology, and Immunology, University of Ottawa, 451 Smyth Road, Ottawa, ON K1H 8M5, Canada.

## Abstract

Mitochondria are the key generators of ATP in a cell. Visually, they are highly dynamic organelles that undergo cellular fission and fusion events in response to changing cellular energy requirements. Mitochondria are now emerging as regulators of mammalian cell motility. Here we show that mitochondria infiltrate the leading edge of NIH3T3 fibroblasts during migration. At the leading edge, we find that mitochondria move to and tether to Focal Adhesions (FA). FA regulate cell migration by coupling the cytoskeleton to the Extracellular Matrix through integrin receptors. Importantly, we find that inhibition of mitochondrial ATP generation concomitantly inhibits FA size. This suggests that mitochondrial energy production regulates migration through FA control.

## Introduction

Mitochondria have important roles in aerobic energy generation [1], cell death [2] and aging [3]. In addition, mitochondria contribute to Ca^2+^ homeostasis, and the metabolism of amino acids, lipid and nucleotides [4, 5]. Dysfunction of mitochondrial metabolism and dynamics contributes to cancer development [6] and a broad spectrum of human diseases [7]. Mitochondria are highly dynamic organelles with fission and fusion events continually reshaping their morphology [8]. Mitochondria vary in size from individual organelles in the submicron length range to large interconnected tubular networks spanning the cytoplasm. It is now recognized that contact between mitochondria and other organelles is an important part of cellular physiology and homeostasis [9, 10]. For example, contact with the Endoplasmic Reticulum licenses mitochondrial fission [11, 12] and mitochondrial derived membranes are part of the peroxisome biogenesis pathway [13].

Mitochondria are emerging as novel regulators of cell motility [14, 15]. Cell migration regulates several important physiological processes, among them embryonic development, tissue morphogenesis and the immune response[16]. Dysregulated migration is often associated with cancer development and metastatic progression [17]. Fragmented mitochondria are found to infiltrate the leading edge of breast and ovarian cancer cell during migration [14, 15]. Inhibition of either mitochondrial ATP generation [14] or Ca^2+^ uptake through silencing of the Mitochondrial Calcium Uniporter (MCU) [15] inhibits motility.

An unresolved question, however, is to what part of the cell migration machinery do mitochondria regulate and what motility structures do they move and tether to. In this report, we find that during fibroblast migration mitochondria move to and tether to Focal Adhesions (FA). FA are multi-protein adhesive structures on the basal cell surface that couple the extracellular matrix (ECM) to intracellular actin fibers via clusters of transmembrane integrin receptors [18, 19]. FA allow for the propagation of mechanical forces within the cell and into the external environment [18, 19]. We find that inhibition of mitochondrial ATP generation decreases FA size, identifying FA as a migratory structure regulated by mitochondrial contact and ATP generation.

## Results & Discussion

### Mitochondria move to the leading edge during migration

In both breast and ovarian cancer cells, mitochondria infiltrate into the leading edge lamellipodia [14, 15, 20]. To determine whether or not this was the case in non-transformed cells, we used live cell microscopy to image mitochondria in freely migrating NIH3T3 mouse fibroblasts (Fig 1a, Supplementary Video 1). In these cells, we observe that as the cell moves forward, multiple mitochondria infiltrate the leading edge. To extend these findings, we tracked individual mitochondria in NIH3T3 cells expressing a fluorescently tagged Cortactin protein (Fig 1b, Supplementary Video 2). Cortactin localises prominently to the lamellipodial edge in migrating cells and regulates cell migration by binding to the Arp 2/3 complex, stabilizing actin branches and activating multiple signaling pathways necessary for motility [21]. As shown in Fig 1b, multiple mitochondria with various starting locations move towards the Cortactin polarized edge (red arrow) during migration (direction indicated by white arrow). In migratory NIH3T3 cells, as in other cell types [22, 23], mitochondria move along microtubules from the cell interior (Fig 1c). However, once in the leading edge, which is thinner than the majority of the cell body, they do not associate with tubulin (Fig 1c). Thus, the presence of mitochondria at the leading edge does not absolutely require microtubule dependent processes.

**Figure 1.**
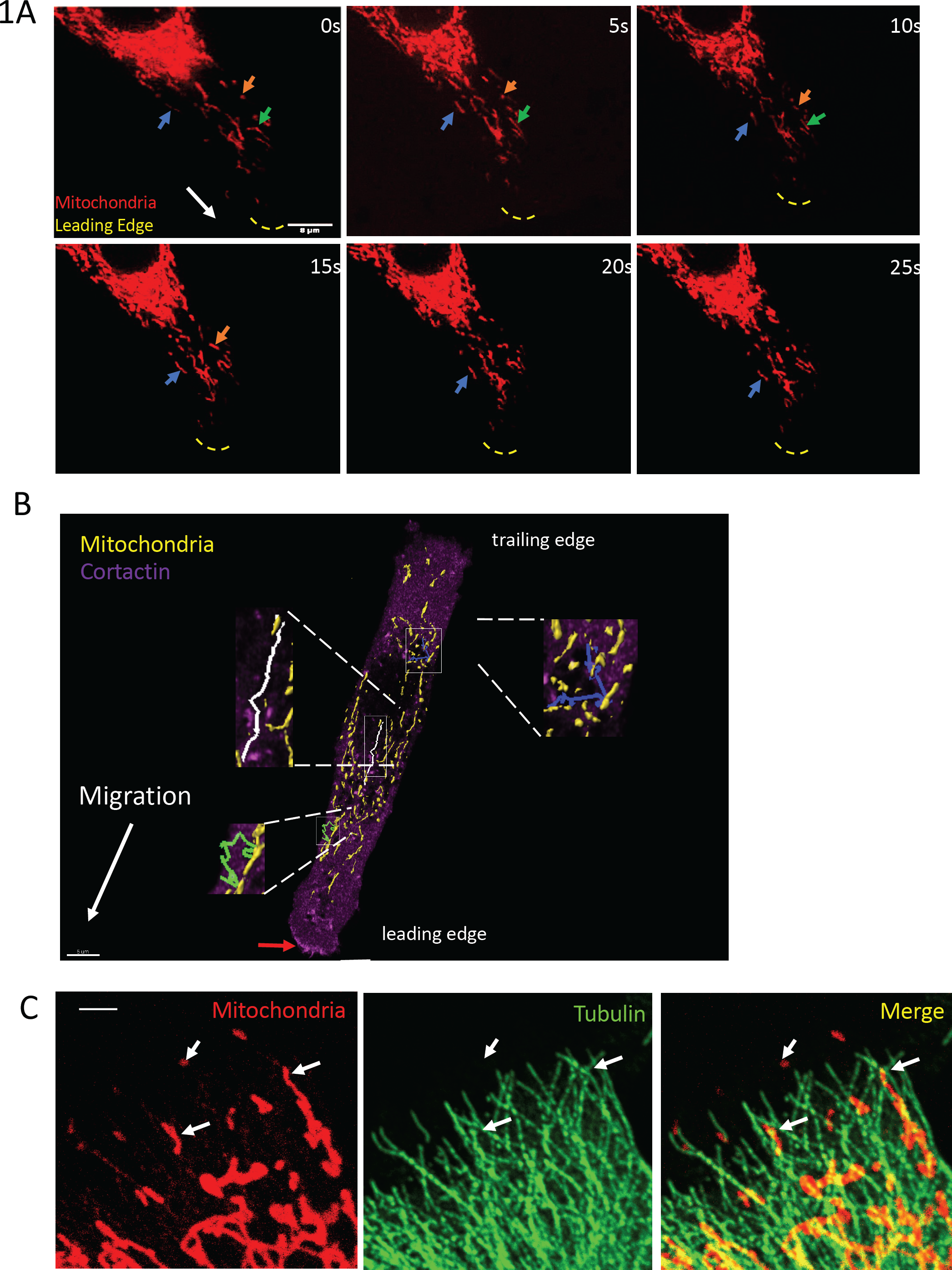
Mitochondria infiltrate the leading-edge during migration. **A)** Mitochondria (colored arrows) move towards the leading edge in migrating NIH3T3 fibroblasts. The direction of migration is indicated by the white arrow. The yellow dashed line indicates the plasma membrane edge from the DIC image (Supplementary Video S1). **B)** Mitochondria move towards the Cortactin edge (red arrow) during cell migration (white arrow) (Supplementary Video S2). **(C)** Mitochondria move along microtubules during migration. Scale bar is 1 μm.

### Mitochondria contact focal adhesions during migration

The infiltration of mitochondria towards the migratory leading edge suggests an important role for them in sustaining cell migration. Previous reports indicate that regulation of calcium flux via the Mitochondrial Calcium Uniporter (MCU) and energy levels via the AMP-activated protein kinase (AMPK) are two pathways through which mitochondria affect migration [14, 15]. However, it is unknown what migratory structures in lamellipodia mitochondria might be interacting with.

We speculated that mitochondria might be interacting with Focal Adhesions (FA) since these adhesive structures position themselves at the periphery of a cell and are frequently found at or close to the lamellipodial edge. To test this idea, we used live cell microscopy to simultaneously visualize mitochondria and the Talin protein during fibroblast migration (Fig 2A and Supplementary Video 3). Talin is a component of FA that links transmembrane integrin receptors to actin fibers in the cytosol [24]. We reasoned that mitochondria might regulate cell migration by interacting with FA since functioning FAs are required for many forms of mammalian cell motility [25]. Fig 2A and Supplementary Video 3 shows the leading edge of a migrating cell where one peripheral mitochondrion undergoes a fission event (f labelled arrow) and the budded mitochondrion sequentially contacts one FA and then another. In fixed cells (Fig 2B), multiple FA in the cell periphery are observed tethered to mitochondria. Surface rendering of the fixed cells (Fig 2C), shows further evidence of FA/mitochondrial tethering. Thus, FA represent a new addition to a growing list of cellular structures that are in physical contact with mitochondria.

**Figure 2.**
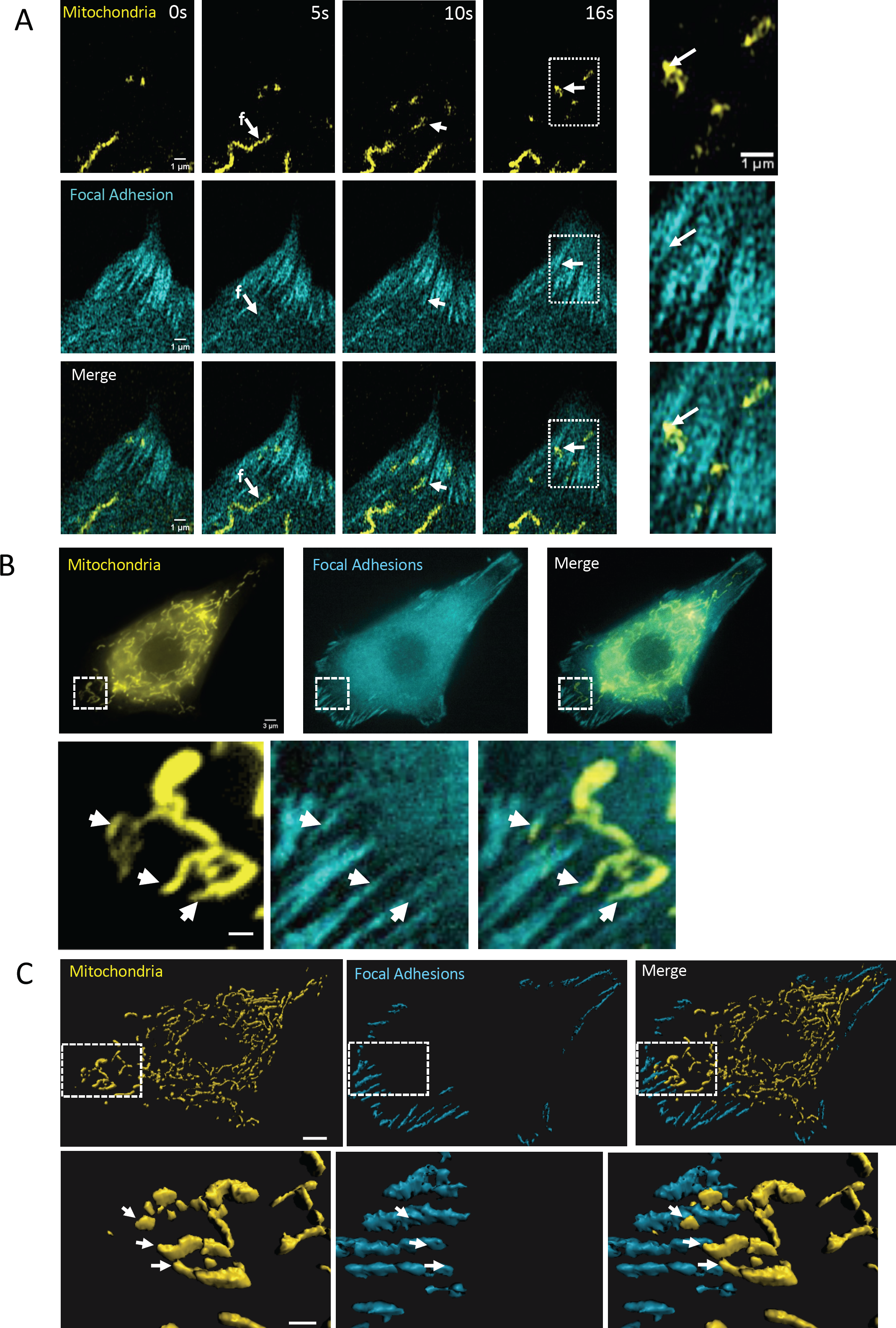
Mitochondria move to and tether to Focal Adhesions During migration. **A)** During migration, a mitochondrion at the leading edge undergoes a fission even (f) and moves to one focal adhesin (Talin) and then another. Inset shows an enlarged image of the boxed rectangle (Supplementary Video 3). **B)** In a fixed cell, mitochondria (MitoTraker) can be seen interacting with Focal Adhesions (Vinculin). Lower images are an enlargement of the boxed rectangle. Scale bar is 1 uM. **C)** Surface reconstruction of the image in (B) shows contact between mitochondria and focal adhesions. Scale bar in the upper and lower image set are 4 and 1 μm respectively.

### Inhibition of mitochondrial activity reduces Focal Adhesion size

We next examined what functional role mitochondria might have on FA structure. First, we used the TMRE dye to measure mitochondrial membrane potential in migrating cells. As shown in Fig 3A, mitochondria tethered to FA have varied mitochondrial membrane potential, some exhibiting high membrane potential while others did not. To determine whether or not mitochondrial activity had any relationship with FA structure, we treated NIH3T3 cells with oligomycin. Oligomycin inhibits mitochondrial ATP generation by preventing ATP synthase activity [26] and has been previously been shown to retard migration of ovarian cancer cells [14]. We treated cells with oligomycin and used an image analysis program [27] to quantitate FA number and size four hours later. As shown in Fig 3B, the mean number of focal adhesions per cell is not affected by oligomycin. Control NIH3T3 cells have an average of 29.8 FA per cell, not statistically different than the 29.2 and 29.8 FA/cell in 1μM and 2μM oligomycin treated cells. On the other hand, oligomycin treated cells significantly have shorter FA compared to control cells, which have an average FA length of 2.78 μm, cells treated with 1 μM or 2 μM oligomycin had an average FA length of 2.33 μm and 2.22 μm respectively. Cumulative Distribution function of the cells shows that oligomycin significantly decreases FA length, consistent with the idea that mitochondrial contact with FA regulates their size (Figure 3c).

**Figure 3.**
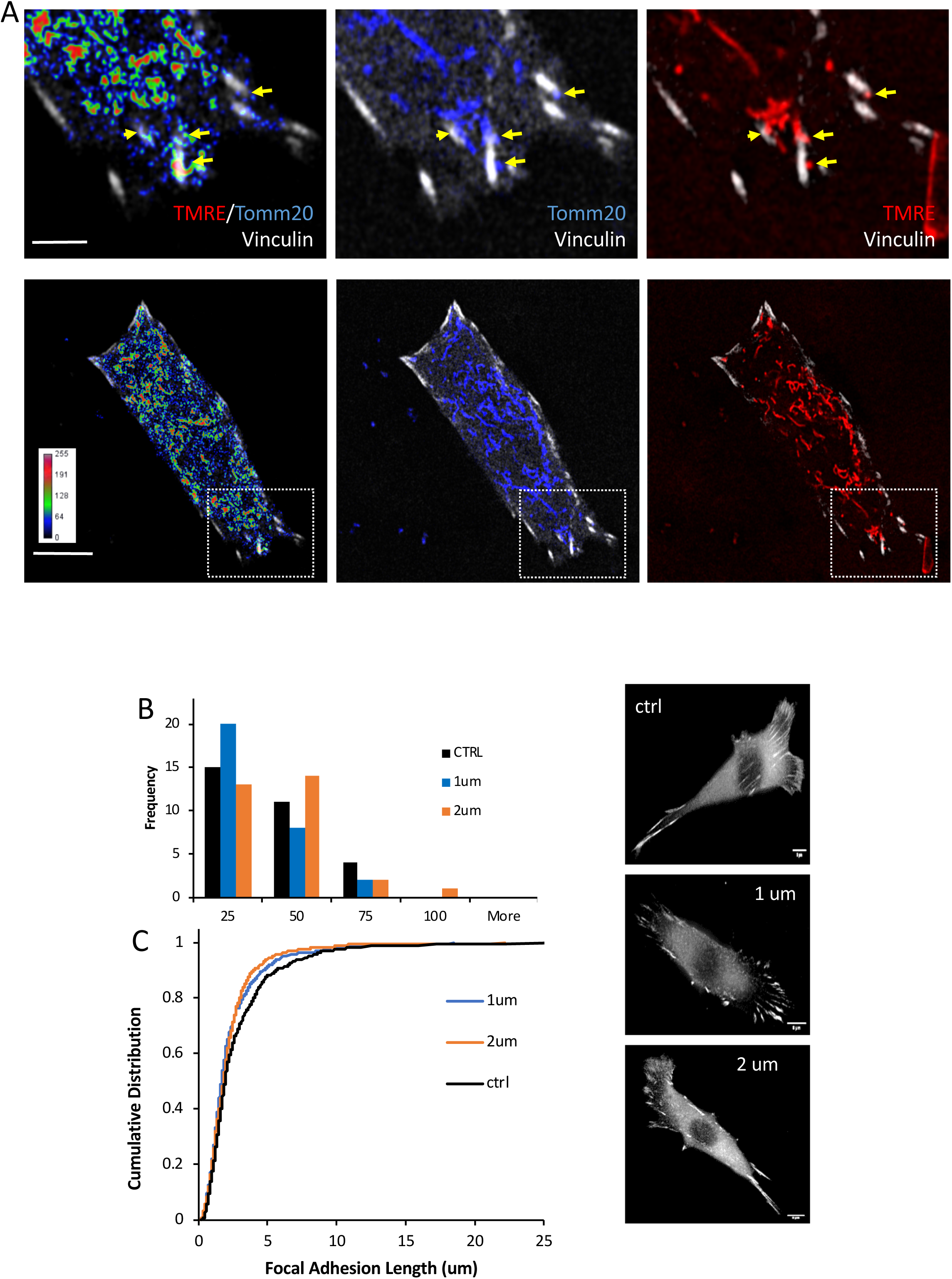
Inhibition of mitochondrial ATP generation reduces focal adhesion number. **A)** TMRE-based measurement of mitochondrial membrane potential (TMRE/Tomm20 ratio). Upper images are an enlarged version of the box in the lower image set. Yellow arrow identify mitochondria in contact with Focal Adhesions. **B)** Distribution of FA number in control cells and those treated with oligomycin. **C)** Cumulative Distribution Function of FA lengths in cells treated with Oligomycin. Focal Adhesion size is derived from three independent experiments each of 30 cells per condition. Representative images are shown in the right panel.

FA are a multi-protein assembly where transmembrane integrin heterodimers binds to the ECM and are linked to the actin cytoskeleton through adapter proteins such as talin, vinculin and paxillin. A mature FA, generally 2 μm wide x 3-10 μm long, is created from smaller adhesive structures described as focal complexes or nascent adhesions [19]. The regulation of FA size by mitochondria suggests that full FA assembly requires mitochondrial action. Our observation that oligomycin decrease FA length is consistent with the idea that localized ATP generation by mitochondria is a key part of FA maturation. High local concentrations of ATP could support actin polymerization or the signaling processes necessary for full FA assembly. FA size has been shown to regulate cell speed, with shorter focal adhesions were associated with slower cell speeds through a biphasic relationship (Kim & Wirtz, 2013). Thus, a requirement for mitochondria in FA maturation provides a plausible explanation for how mitochondrial metabolism might regulate motility.

Interaction between mitochondria and other cellular structures is now understood to be an important aspect of the homeostasis of multiple organelles [9]. Mitochondria make important functional contacts with the endoplasmic reticulum (ER), vacuoles and peroxisomes [9, 13]. Mitochondrial contact with the ER regulates mitochondrial division [12] and also allows ER-derived lipids to move to mitochondria. With respect to non-membrane bound structures, mitochondria are dependent on microtubules for their transport and Kinesin and Dynein motor proteins link mitochondria to tubulin. Our work shows that Focal Adhesions are among the list of organelles and cellular structures interacting with mitochondria and is consistent with the idea that mitochondria regulate cell migration, at least in part, by regulating cellular adhesion.

## ACKNOWLEDGEMENTS

The authors thank Skye McBride and Chloe van Oostende for training and assistance with microscopy. We thank John Copeland, Daniel Jacobsen, Mireille Khacho and Julie St-Pierre for helpful discussion. This work is supported by an operating grant from NSERC (JML).

## METHODS

### Cell Culture

NIH3T3 cells were obtained from the ATCC. Cells were cultured at 37°C, 5% CO_2_ in DMEM (Thermo Scientific, Burlington, Canada)supplemented with 10% FBS (Thermo Scientific, Burlington, Canada), 1mmol/L sodium pyruvate (Thermo Scientific), and penicillin-streptomycin.

### Live cell imaging

For imaging, cells were trypsinized from 80-100% confluent plates and seeded at 10^5^ cells per 35mm µ-Dish high-wall (Ibidi) or in a 6-well plates (Corning) containing High Precision 1.5H cover slips (Deckglaser) and left to adhere to the surfaces overnight. The next day cells are transfected using Lipofectamine 2000 (11668-019) from Thermo Fisher at a 1:1.5 Lipo:DNA ratio for GFP-Cortactin (Addgene 50728), mEmerald-Talin (Addgene 54266) or mCerulean3-Tomm20 (Addgene55450). 1000 ng of cortactin and of talin was used for respective transfections. Cells were imaged in Gibco phenol-free DMEM and mitochondria were stained with Thermo Fisher Mitotracker Red CMXRos (M7512) for 30 minutes. Live cell cortactin images are taken and deconvolved on the DeltaVision Elite-Olympus IX-71 with FemtoJet Microinjector using version 7 of Softworx. Deconvolution is done with 5 iterations. The Zeiss LSM880 AxioObserverZ1 inverted confocal microscope with AiryScan FAST mode at 63X objective was used for live-cell imaging of talin images. For mitochondrial activity experiments, transfected cells were incubated to a final concentration of 100mM TMRE (ThermoFisher Scientific #T669) for 15 minutes at 37°C. The TMRE potential was detected upon excitation with the 561nm laser line and mCerulean3 detected upon excitation with the 405nm laser line.

### Fixed Cell Microscopy

Cells grown on 18mm coverslips were incubated for 15 min at 37°C with a Mitotracker™Red CMXRos to a final concentration of 100-200 nM. Cells were fixed with 4% PFA/PBS for 10 min at room temperature, washed 3x with PBS and permeabilized with 0.1% Triton X-100/PBS for 15 min. Cells were then blocked for 1 hr at room temperature or overnight at 4° with 3% FBS/0.1% Triton X-100 in PBS. Cells were then stained for 1 hr at room temperature with Monoclonal Anti-Vinculin antibody (Sigma V9131). Cells were washed with 3X PBS. Cells were then stained for 1 hr (RT) with secondary antibody, Alexa FluorTM Plus 488 (A32723, Thermofisher). Cells were then washed 3X with PBS. Cells were imaged on the Zeiss AxioObserver Z1 microscope using a 63X, 1.4 NA objective and deconvolved using Zen blue edition under fast iterative parameters and bad pixel correction. Spatial reconstruction of images were generated using surface rendering on Imaris software. The quantitation of focal adhesions immunostained for vinculin was performed using ImageJ.

## ADDITIONAL INFORMATION

**Supplementary video SV1.** Migrating NIH 3T3 fibroblast with mitochondria visualized with MitoTacker Red and the cell body visualized with DIC.

**Supplementary video SV2.** Migrating NIH 3T3 fibroblast transfected with GFP-Cortactin and with mitochondria visualized with MitoTacker Red. Cortactin has been pseudo-coloured purple and mitochondria pseudo-coloured yellow for visual clarity.

**Supplementary video SV2.** Migrating NIH 3T3 fibroblast transfected with GFP-Vinculin and with mitochondria visualized with MitoTacker Red. Vinculin has been pseudo-coloured cyan and mitochondria pseudo-coloured yellow for visual clarity.

## Notes

**Funding:** This work was supported by operating funds from the Natural Sciences and Engineering Research Council of Canada (JML).

